# Cognitive strategy accounts for failure on a hippocampal relational memory task in non-human primates

**DOI:** 10.1101/2025.04.23.650325

**Authors:** Casey R. Vanderlip, Shelby R. Dunn, Payton A. Asch, Courtney Glavis-Bloom

## Abstract

Relational memory, the ability to flexibly encode and retrieve associations among distinct elements, is critically dependent on the hippocampus and declines with age in humans. The Transverse Patterning (TP) task is designed to probe relational memory by requiring learning of hierarchical, circular stimulus relationships (e.g., A+ B-, B+ C-, C+ A-), a structure akin to rock-paper-scissors. In humans, TP performance is reliably impaired by hippocampal damage and aging. In non-human primates, however, findings have been inconsistent with some studies demonstrating clear hippocampal dependence, while others report no impairment, or even improvements, following hippocampal lesions. This raises the possibility that species differences in cognitive strategy use may underlie these divergent outcomes. We hypothesized that non-human primates rely on an elemental learning strategy, supported by corticostriatal systems, even when relational memory is required. To test this, we trained young and aged common marmosets (*Callithrix jacchus*) on the TP task and several control tasks designed to isolate elemental versus configural learning. Marmosets successfully acquired reward contingencies for individual stimulus pairs but failed when success required integrating all three stimulus relationships. In contrast, all animals readily acquired control tasks solvable via simple stimulus-response associations. Notably, there was no evidence of age-related impairment on TP or control task performance. Given the early vulnerability of the hippocampus to aging and the relative preservation of striatal systems, this pattern further supports the conclusion that marmosets rely on a habit-based learning strategy that is poorly suited to relational demands. These findings suggest that humans and non-human primates may approach the same tasks using different cognitive strategies. This has critical implications for interpreting cross-species differences in memory performance and highlights the need to validate which neural systems a task engages in each species before using it as a translational model of hippocampal function or cognitive aging.

## INTRODUCTION

Relational memory refers to the ability to flexibly encode and retrieve associations among elements that are not inherently linked (Cohen et al., 1999; Davachi & Wagner, 2002; Ryan et al., 2000). This capacity enables the flexible integration of experiences across time and context and is considered foundational to episodic memory. Converging evidence from behavioral, neuroimaging, and lesion studies implicates the hippocampus as essential for relational memory (Banta Lavenex et al., 2006; Cohen et al., 1999; Giovanello et al., 2009; Ryan et al., 2000). Importantly, some theories propose that relational memory is not merely one of many functions of the hippocampus - it is the fundamental process that defines its role in cognition (Cohen & Eichenbaum, 1993; Konkel, 2009; Kumaran & Maguire, 2005; Ryan et al., 2000).

The Transverse Patterning (TP) task is a classic cognitive paradigm for assessing relational memory. Successful completion of TP requires subjects to learn a set of overlapping stimulus relationships in which no single stimulus is always correct or incorrect. In this task, three stimuli (A, B, and C) are presented in pairs: A is rewarded over B, B over C, and C over A – a structure like the game rock-paper-scissors (Astur & Sutherland, 1998; Reed & Squire, 1999; Rickard et al., 2006). Because the correct choice depends on the *relationship between* stimuli rather than any individual cue, TP cannot be solved through simple stimulus-response associations. Instead, successful performance requires encoding the relative relationships among all stimuli, a hallmark of relational memory. As such, the task is considered to rely heavily on hippocampal function (Rickard et al., 2006).

In humans, performance on the TP task is critically dependent on the hippocampus. Individuals with hippocampal damage consistently fail to acquire the task structure, despite intact performance on simpler learning paradigms that rely on direct stimulus-response associations (Astur & Sutherland, 1998; Reed & Squire, 1999; Rickard et al., 2006; Rickard & Grafman, 1998). TP performance also declines with age, mirroring age-related changes in hippocampal structure and function (Gracian et al., 2016; Leirer et al., 2010; Meltzer et al., 2009). In contrast, elemental learning, which involves forming independent associations between stimuli and outcomes and is mediated by striatal circuits, remains relatively preserved across the lifespan (Cox et al., 2008; Gardner et al., 2020; Hill et al., 2023). This dissociation reinforces the value of the TP task in isolating hippocampal-dependent relational memory from striatal, habit-based learning mechanisms (Gracian et al., 2016; Rickard & Grafman, 1998).

In non-human primates, however, the role of the hippocampus in TP performance is far less consistent. Some studies report robust impairments following hippocampal lesions, aligning with findings in humans (Alvarado & Bachevalier, 2005). Yet others show no effect of lesions, or even facilitation of learning (Saksida et al., 2006), suggesting that the hippocampus is not critical for this task. These contradictory outcomes raise the possibility that different species rely on distinct cognitive strategies when performing the TP task. Unlike humans, macaque monkeys often require hundreds or even thousands of trials to reach criterion, suggesting they may be solving the task through incremental stimulus-response learning rather than through configural (Alvarado & Bachevalier, 2005; Bussey et al., 1998; Saksida et al., 2006). Moreover, while TP performance declines reliably with age in humans, no studies have examined age-related effects in non-human primates, leaving open questions about how aging affects relational memory across primate species.

Together, these findings challenge the assumption that TP engages the same cognitive and neural mechanisms across species and underscore the need to consider species-specific strategies underlying task performance. We hypothesized that non-human primates rely on habit-based strategies to solve two-alternative forced-choice tasks, utilizing elemental learning supported by striatal and cortical systems. This approach is effective for many commonly used tasks, which can be solved through reinforcement alone, without requiring relational memory. In contrast, TP presents a unique cognitive demand: because the correct choice response depends on the relative relationships among stimuli, rather than on the stimuli in isolation, success requires a configural learning strategy supported by the hippocampus. However, this raises an important question: in tasks like transverse patterning that *cannot* be successfully solved using a habit-based approach, will non-human primates continue to rely on suboptimal strategies, or will they recruit relational memory systems when needed? Understanding whether this species difference in strategy use is flexible, or rigid is critical for interpreting cross-species findings and for assessing the translational validity of tasks intended to probe hippocampal function.

To address this, we tested common marmosets (*Callithrix jacchus*) on the TP task to determine whether they would utilize a default habit-based strategy or engage hippocampal-dependent relational memory. The marmoset is an increasingly important model in cognitive neuroscience, offering unique advantages for translational aging research due to its shared neuroanatomical and functional brain organization with humans, complex social and cognitive repertoire, and relatively short lifespan (Glavis-Bloom et al., 2023; Vanderlip et al., 2025). By examining how marmosets approach a task that cannot be solved through elemental learning alone, we aimed to assess whether non-human primates flexibly adopt configural strategies when the task demands it.

We found that marmosets successfully learned the reward contingencies for individual problems but failed to acquire the full TP structure once success depended on understanding the relationships among all three stimuli, suggesting a failure to engage relational memory processes. In contrast, performance on control tasks that could be solved through simple stimulus-response associations was intact, indicating that the deficit was not due to a general learning impairment. By testing a cohort of marmosets spanning the full adult lifespan, we were also able to assess age-related effects on TP performance. Strikingly, we found no evidence of age-related decline, in contrast to the robust age effects observed in humans. Given that the hippocampus is one of the first regions to show age-related decline, while striatal learning systems remain largely intact, these findings further support the idea that marmosets rely on habit-based strategies to solve the task. Together, our results highlight species differences in cognitive strategy use and underscore the importance of understanding not just *what* animals learn, but *how* they learn it.

## METHODS

### Subjects

Twelve common marmosets (*Callithrix jacchus*; 7 males, 5 females) between the ages of 4.5 and 13 years participated in the study (Table 1). Animals were categorized as Young if they were younger than 8 years, and as Aged if they were 8 years or older. While there is currently no consensus on the exact age at which marmosets are considered aged, they are commonly classified as aged between 7 and 8 years, as some age-related pathologies (e.g., reduced neurogenesis) begin to emerge around this time (Perez-Cruz & Rodriguez-Callejas, 2023; Tardif et al., 2011). All marmosets had prior experience with touchscreen-based cognitive testing in their home cages (Glavis-Bloom et al., 2022; Vanderlip et al., 2024). Animals were housed either singly or in pairs, with environmental enrichment including hammocks and manzanita branches. For testing, pair-housed animals were temporarily separated by a divider in the home cage. All animals maintained visual and auditory access to other marmosets at all times. All experimental procedures were approved by the Salk Institute Institutional Animal Care and Use Committee and followed the guidelines of the NIH Guide for the Care and Use of Laboratory Animals.

**Table 1.** Description of subjects.

| Marmoset ID | Age | Sex |
| --- | --- | --- |
| FI | 4.5 | F |
| WA | 5.3 | M |
| AN | 5.6 | F |
| BL | 7.3 | M |
| JA | 7.7 | M |
| LA | 7.9 | M |
| VE | 8.2 | F |
| CC | 9.6 | F |
| CP | 10.1 | F |
| PE | 10.1 | M |
| TR | 10.9 | M |
| CI | 13.0 | M |

### Equipment and Software

Marmoset home cages were custom-designed to include a testing chamber located in the upper corner of each enclosure, where a 10.4-inch infrared touchscreen station (Lafayette Instrument Company, Lafayette, IN) was mounted. Access to the testing chamber was restricted outside of testing sessions. During testing, animals could freely enter and exit the chamber through a small doorway. Liquid rewards (e.g., apple juice) were delivered into a small basin beneath the touchscreen upon correct responses. Stimulus locations on the screen were indicated by cutouts in a plastic overlay placed in front of the display, providing a consistent visual frame for response options.

All tasks were programmed and controlled using Animal Behavior Environment Test (ABET) Cognition software (Lafayette Instrument Company, Lafayette, IN). The software recorded trial-level data including response accuracy, location of stimuli and responses on the screen, and response latencies. Stimuli consisted of simple black and white geometric shapes and lines presented on a black background.

### Cognitive Testing

Marmosets were tested for 1 to 3 hours per day, 3 to 5 days per week. They worked for fluid rewards and were not subjected to food or water restriction. Marmosets were tested on a Transverse Patterning (TP) task, and on three control tasks designed to isolate different cognitive demands. All tasks used a two-alternative forced-choice design. At the start of each trial, marmosets were required to initiate the trial by touching a centrally located trial initiation stimulus. Following this, two visually distinct black and white stimuli were displayed on the touchscreen, with left/right position counterbalanced across trials. Selecting the correct stimulus resulted in immediate delivery of a liquid reward. Incorrect responses or omissions (no response within 12 seconds) triggered a 5-second timeout with a blank screen. All trials, regardless of outcome, were followed by a 5-second inter-trial interval. Unique stimuli were used for every problem and condition.

### Cognitive Task Descriptions

Depictions of each of the cognitive tasks can be found in Table 2.

**Table 2.**
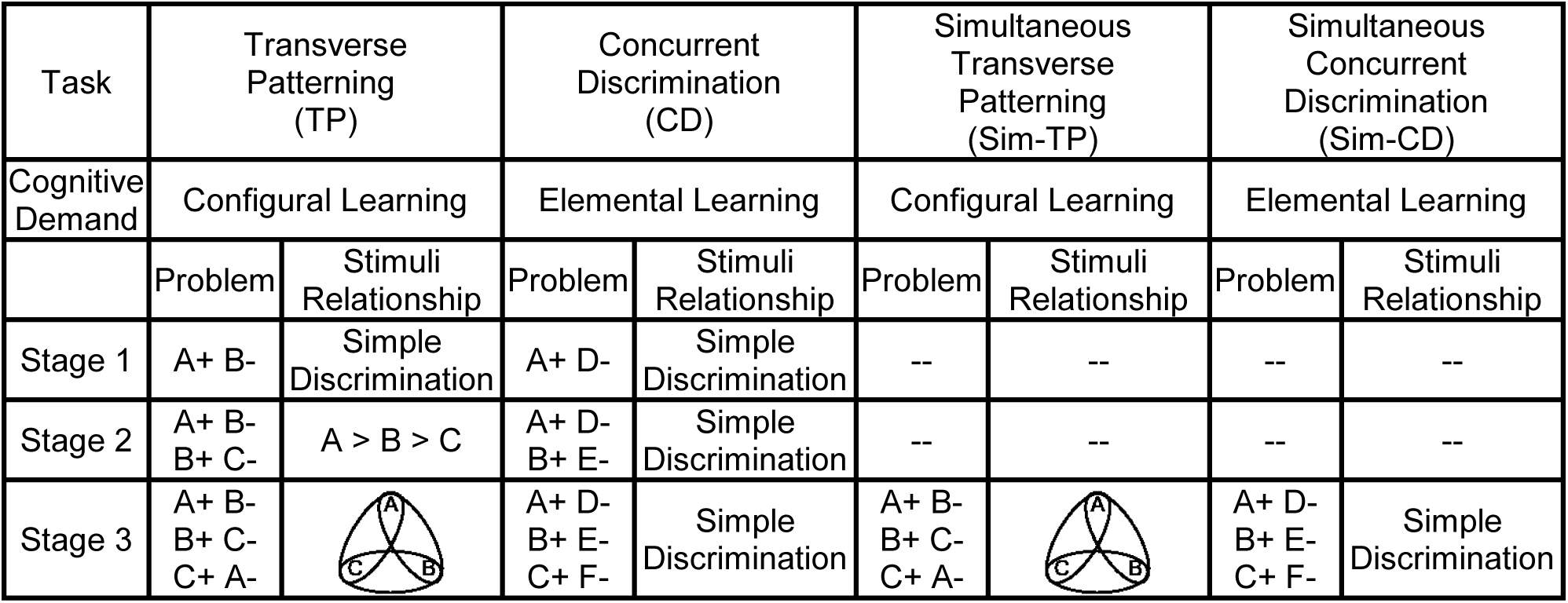
Cognitive task depictions.

| Task | Transverse Patterning (TP) |  | Concurrent Discrimination (CD) |  | Simultaneous Transverse Patterning (Sim-TP) |  | Simultaneous Concurrent Discrimination (Sim-CD) |  |
| --- | --- | --- | --- | --- | --- | --- | --- | --- |
| Cognitive Demand | Configural Learning |  | Elemental Learning |  | Configural Learning |  | Elemental Learning |  |
|  | Problem | Stimuli Relationship | Problem | Stimuli Relationship | Problem | Stimuli Relationship | Problem | Stimuli Relationship |
| Stage 1 | A+ B- | Simple Discrimination | A+ D- | Simple Discrimination | -- | -- | -- | -- |
| Stage 2 | A+ B-<br>B+ C- | A > B > C | A+ D-<br>B+ E- | Simple Discrimination | -- | -- | -- | -- |
| Stage 3 | A+ B-<br>B+ C-<br>C+ A- |  | A+ D-<br>B+ E-<br>C+ F- | Simple Discrimination | A+ B-<br>B+ C-<br>C+ A- |  | A+ D-<br>B+ E-<br>C+ F- | Simple Discrimination |

### Transverse Patterning (TP)

The TP task involved three overlapping discriminations: A+ B-, B+ C-, and C+ A-, requiring subjects to learn the relative relationships among stimuli rather than relying on simple reward history. Problems were introduced sequentially over three stages. In Stage 1, animals were trained only on Problem 1 (A+ B-). In Stage 2, Problem 2 (B+ C-) was added alongside continued presentation of Problem 1 such that approximately half of the trials were of each Problem. In Stage 3, all three problems were presented in random order. Successful performance on this final stage requires relational learning rather than simple stimulus– response associations, as no single stimulus is consistently correct. To preserve motivation for future cognitive testing, animals were removed from the task if performance on all three problems declined to chance levels for a sustained period during Stage 3.

### Concurrent Discrimination (CD)

The CD control task assessed whether marmosets could successfully acquire and retain multiple stimulus-response associations when those associations did not involve relational or overlapping elements. Like the TP task, CD was structured across three sequential stages, but the stimulus pairs were non-overlapping (e.g., A+ D-, B+ E-, C+ F-). Because each problem could be learned independently, the CD task could be solved using an elemental learning strategy without engaging the configural processing or interference resolution typically supported by the hippocampus. Successful performance on this task would indicate that any deficits observed on TP were not due to general limitations in memory capacity or the ability to manage multiple associations across time.

### Simultaneous Transverse Patterning (Sim-TP)

The Sim-TP control task was designed to test whether extended exposure to individual discriminations (e.g., A+ B- and B+ C-) in the TP task promoted the use of simple, non-relational rules that interfered with acquiring the full relational structure. In this variant, all three overlapping problems (A > B, B+ C-, C+ A-) were introduced simultaneously from the start, preventing animals from relying on simple, non-relational rules learned through incremental stimulus-response learning on earlier problems. Like the original TP task, no stimulus was consistently rewarded across all trials, preserving the requirement for configural learning.

### Simultaneous Concurrent Discrimination (Sim-CD)

The purpose of the Sim-CD control task was to determine whether poor performance on the Sim-TP task reflected a general difficulty with learning multiple discriminations simultaneously, or a specific impairment in acquiring the relational structure across overlapping stimuli. Strong performance on this control task would indicate that marmosets can manage multiple discriminations in parallel when relational memory is not required, supporting the interpretation that TP deficits reflect a specific failure to engage configural learning strategies. Therefore, this task was designed to match the cognitive load and structure of the Sim-TP task while eliminating the relational demands. Animals were presented with three non-overlapping problems from the start: A+ D-, B+ E-, C+ F-, and each problem had a consistent correct choice. Because there were no shared stimuli across problems, the task could be solved using elemental learning strategies without requiring relational processing.

In all tasks, each stage continued until animals reached a criterion of at least 9 correct responses out of 10 consecutive trials for each problem in the stage. Trial order was pseudorandomized, and stimulus positions were counterbalanced across trials to control for side biases. Marmosets remained in each stage until they reached the criterion or until performance on each stimulus converged toward chance, a decision made to avoid extinguishing touch screen behavior.

### Statistical Analyses

All statistical analyses were conducted in R (R Core Team, 2023). Because the number of trials needed to reach criterion varied across animals, stages, and tasks, we standardized performance trajectories by expressing trial progression as a proportion of total trials completed within each stage. This allowed us to compare learning dynamics across animals and conditions on a common timescale. We used two complementary approaches to assess performance over time. First, we applied locally estimated scatterplot smoothing (LOESS) to accuracy data across trials within each stage. LOESS is a non-parametric technique that models local trends in the data without assuming a specific functional form, making it well-suited for visualizing learning dynamics. Second, we binned trials into deciles (i.e., 10% segments of total trials per stage) within each stage to assess how accuracy changed over time and across Problems. Previous studies in humans have quantified errors or trials to the performance criterion as the dependent measure (Gracian et al., 2016; Reed & Squire, 1999). Here, since criterion was not met for a subset of tasks, we report accuracy as our primary dependent measure. For comparisons involving two factors, such as Problem number and Trial Bin, we used two-way ANOVAs, followed by Tukey’s HSD post hoc tests to evaluate specific group differences while controlling for multiple comparisons. To examine associations between continuous variables such as age and task performance, we used robust linear regression, which provides more reliable estimates in the presence of outliers or non-normal residuals, both common in studies with small sample sizes. All reported p-values reflect two-tailed tests, with a significance threshold of α = 0.05.

## RESULTS

### Experiment 1

#### Marmosets fail to acquire the relational structure of the Transverse Patterning (TP) task

We trained 12 marmosets on the Transverse Patterning (TP) task across three sequential stages. All animals successfully completed Stage 1 (A+ B-) and Stage 2 (A+ B- and B+ C-), but none were able to learn Stage 3, which required integrating all three overlapping stimulus relationships (A+ B-, B+ C-, C+ A-). Trials for each animal to reach the learning criterion for Stages 1 and 2, and trials until performance converged to chance in Stage 3 are provided in Table 3. To examine learning trajectories, we normalized trials within each stage and applied LOESS smoothing to visualize accuracy over time (Fig 1A).

**Table 3.**
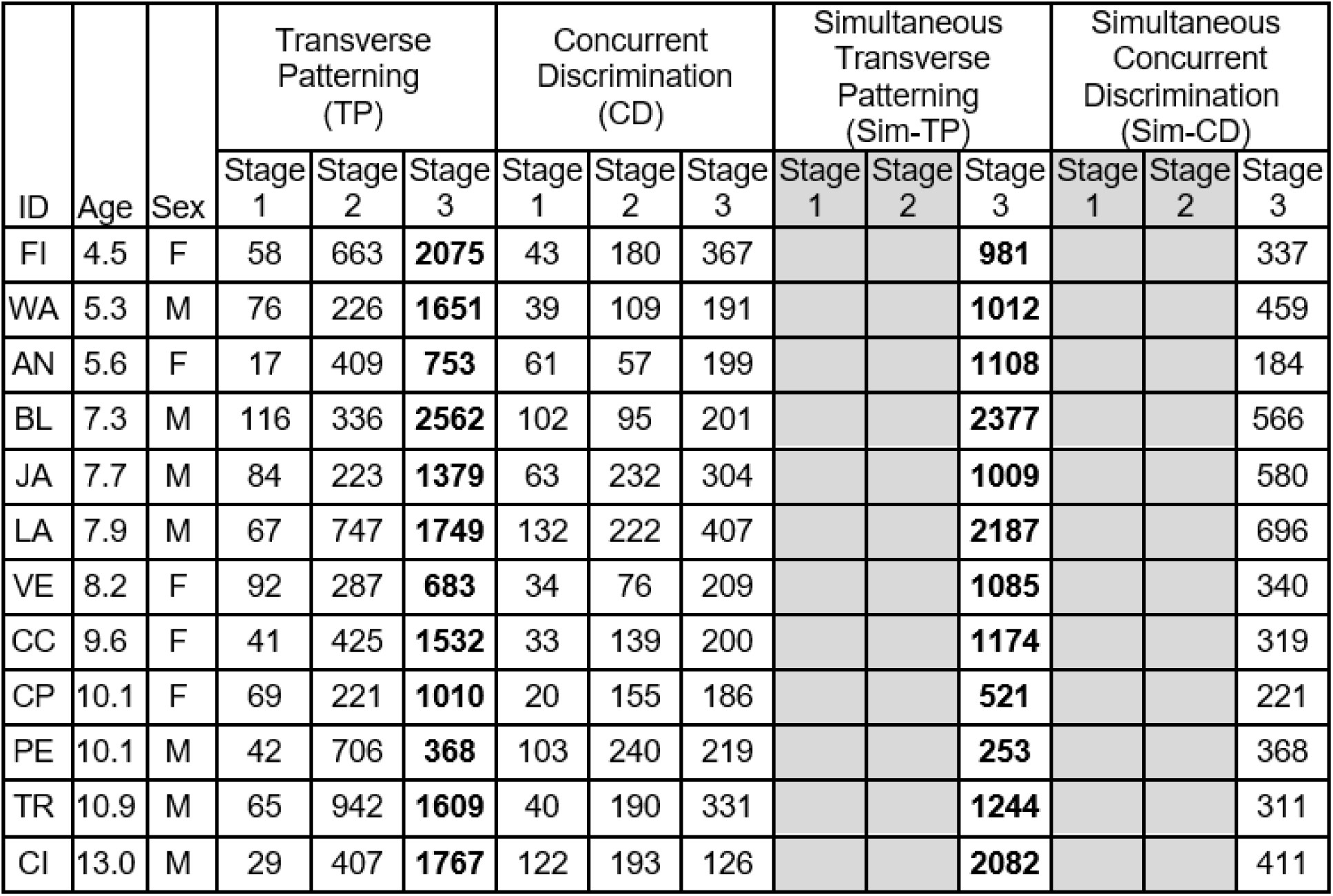
Number of trials completed by each animal for each Stage of each Task. Trial numbers in normal font indicate criterion was reached. Trial numbers in bold font indicate total trials completed before accuracy on each Problem converged at approximately chance levels and testing was terminated.

| ID | Age | Sex | Transverse Patterning (TP) |  |  | Concurrent Discrimination (CD) |  |  | Simultaneous Transverse Patterning (Sim-TP) |  |  | Simultaneous Concurrent Discrimination (Sim-CD) |  |  |
| --- | --- | --- | --- | --- | --- | --- | --- | --- | --- | --- | --- | --- | --- | --- |
|  |  |  | Stage 1 | Stage 2 | Stage 3 | Stage 1 | Stage 2 | Stage 3 | Stage 1 | Stage 2 | Stage 3 | Stage 1 | Stage 2 | Stage 3 |
| FI | 4.5 | F | 58 | 663 | **2075** | 43 | 180 | 367 |  |  | **981** |  |  | 337 |
| WA | 5.3 | M | 76 | 226 | **1651** | 39 | 109 | 191 |  |  | **1012** |  |  | 459 |
| AN | 5.6 | F | 17 | 409 | **753** | 61 | 57 | 199 |  |  | **1108** |  |  | 184 |
| BL | 7.3 | M | 116 | 336 | **2562** | 102 | 95 | 201 |  |  | **2377** |  |  | 566 |
| JA | 7.7 | M | 84 | 223 | **1379** | 63 | 232 | 304 |  |  | **1009** |  |  | 580 |
| LA | 7.9 | M | 67 | 747 | **1749** | 132 | 222 | 407 |  |  | **2187** |  |  | 696 |
| VE | 8.2 | F | 92 | 287 | **683** | 34 | 76 | 209 |  |  | **1085** |  |  | 340 |
| CC | 9.6 | F | 41 | 425 | **1532** | 33 | 139 | 200 |  |  | **1174** |  |  | 319 |
| CP | 10.1 | F | 69 | 221 | **1010** | 20 | 155 | 186 |  |  | **521** |  |  | 221 |
| PE | 10.1 | M | 42 | 706 | **368** | 103 | 240 | 219 |  |  | **253** |  |  | 368 |
| TR | 10.9 | M | 65 | 942 | **1609** | 40 | 190 | 331 |  |  | **1244** |  |  | 311 |
| CI | 13.0 | M | 29 | 407 | **1767** | 122 | 193 | 126 |  |  | **2082** |  |  | 411 |

**Figure 1.**
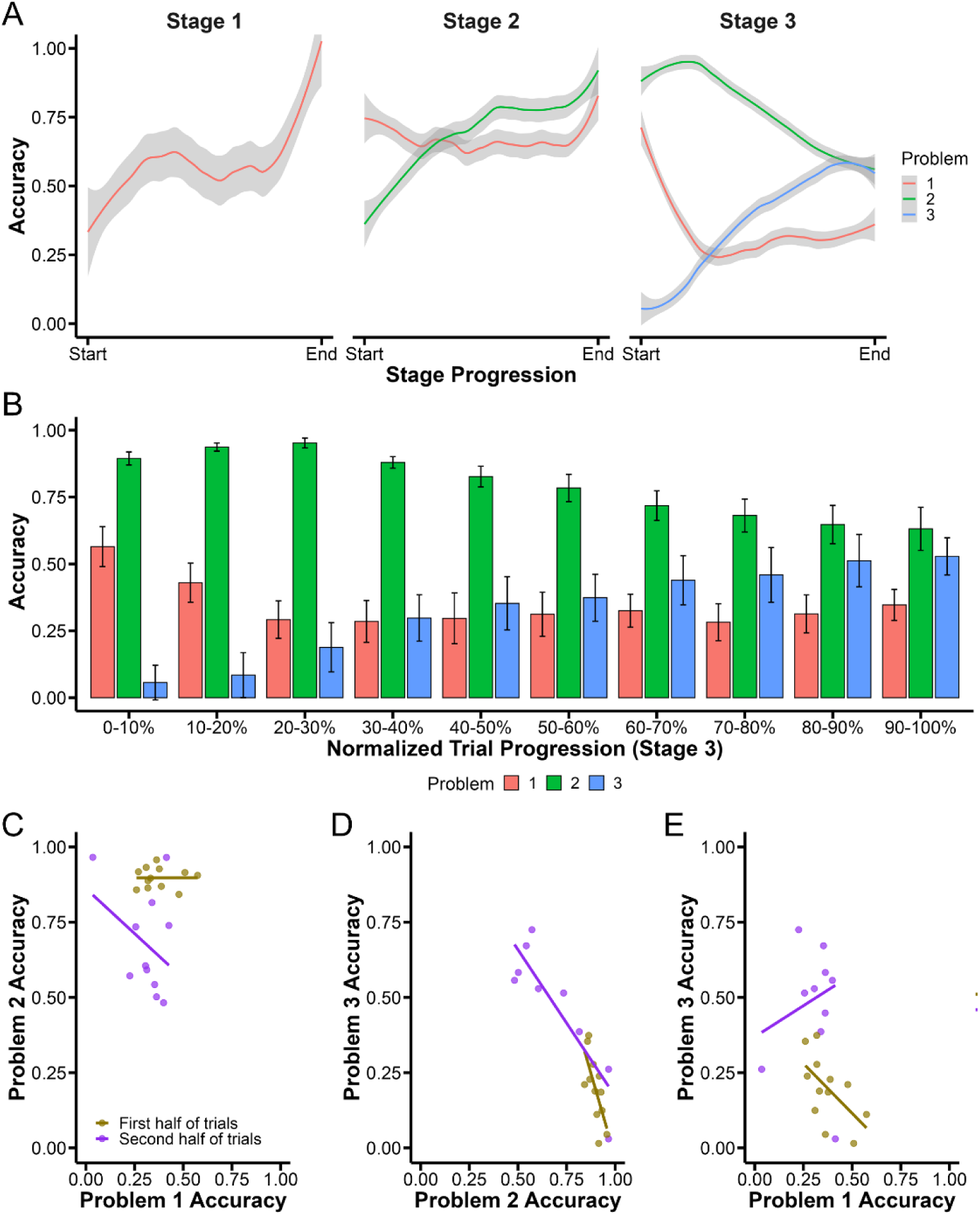
Marmosets fail to acquire the full relational structure of the Transverse Patterning (TP) task. (A) LOESS-smoothed learning curves for each problem across the three stages of TP. Marmosets improved on Problems 1 and 2 in earlier stages but failed to reach criterion in Stage 3, with performance on all three problems converging near chance. (B) Binned accuracy during Stage 3, showing accuracy for each problem across 10% increments of total trials. As Problem 3 accuracy increased, accuracy on Problems 1 and 2 declined, consistent with interference across overlapping associations. (C-E) Scatterplots showing relationships between accuracy on individual problems during Stage 3. (C) Problem 1 vs Problem 2 accuracy; (D) Problem 2 vs Problem 3 accuracy; (E) Problem 1 vs Problem 3 accuracy. Data are separated by trial half (first half in brown, second half in purple). Regression lines show shifting patterns of interference over time. In early trials, performance on Problems 2 and 3 in panel D and Problems 1 and 3 in panel E was negatively correlated, indicating conflict between overlapping stimulus associations. This pattern persisted into the second half of the stage, consistent with a failure to integrate across problems using a configural learning strategy.

During Stage 1, marmosets began at chance and quickly improved, reaching criterion on Problem 1 (A > B). In Stage 2, they acquired the newly introduced Problem 2 (B > C) while maintaining high accuracy on Problem 1, suggesting that learning a second overlapping Problem did not interfere with memory retention of the first. In sharp contrast, Stage 3 was marked by a dramatic shift in performance: accuracy on Problem 1 dropped rapidly below chance, Problem 2 accuracy initially remained high, but then also declined, and accuracy on Problem 3 (C+ A-) showed only modest improvement. Over time, accuracy across all three problems converged around chance. This pattern was observed in every individual marmoset, indicating a consistent and systematic failure to acquire the full relational TP structure.

To quantify these patterns, we binned trials within Stage 3 into deciles and averaged accuracy across animals (Fig 1B). A two-way ANOVA revealed a significant main effect of Problem (*F*(2, 330) = 335.22, *p* < 0.0001) and a significant Problem × Trial Bin interaction (*F*(18, 330) = 12.20, *p* < 0.0001), indicating that performance differed across the three Problems and that these differences changed over time. There was no main effect of Trial Bin alone (*p* = 0.999), indicating that learning did not progress uniformly across problems.

Post hoc comparisons revealed that early in Stage 3 (0-10% of trials), performance was highest on Problem 2, followed by Problem 1, with Problem 3 lowest (all *p*s < .0001). This pattern persisted through the 10–20% bin, with robust differences between all problems (all *p*s < .0001). By the 20-30% bin, performance on Problem 3 had improved, and no longer differed from Problem 1 (*p* = 0.25). From 30-70%, performance on Problems 1 and 3 remained statistically indistinguishable (*p*s > 0.18), while performance on Problem 2 remained significantly higher (all *p*s < 0.0001). In the 7-80% bin, accuracy on Problem 3 surpassed Problem 1 (*p* = 0.018), and this pattern persisted in the 80-90% bin (*p*s < 0.01). By the final decile (9-100%), performance on Problem 1 remained significantly worse than Problems 2 and 3 (*p*s < 0.05), which no longer differed from each other (*p* = 0.25).

These results reveal a progressive pattern of retroactive interference, in which earlier-learned Problems degraded as new, overlapping associations were introduced. Rather than building an integrated relational structure using a configural learning strategy, marmosets appeared to abandon previously learned associations as conflict accumulated.

We hypothesized that marmosets failed to solve the full TP structure because they relied on a stimulus-response association strategy, treating each stimulus (A, B, or C) as independently rewarded or unrewarded based on recent reinforcement history. This elemental learning approach can succeed when problems are learned in isolation but breaks down when stimuli appear in multiple pairings with conflicting outcomes, as required by the relational logic of Stage 3.

To test whether interference emerged between specific problems, we examined correlations in accuracy across problems during the first and second halves of Stage 3. In the first half (0-50% of trials), performance on Problem 1 (A+ B-) and Problem 2 (B+ C-) was not significantly correlated (Fig 1C, β = 0.001, *p* = 0.994), likely reflecting the continued stability of these associations, which had been well learned in the earlier stages. In contrast, performance on Problem 3 (C+ A-) was negatively correlated with both Problem 1 (Fig 1E, β = -0.660, *p* = 0.012) and Problem 2 (Fig 1D, β β = -2.241, *p* = 0.003), suggesting that learning the new C vs A association disrupted performance on previously acquired problems.

This pattern likely reflects conflicting demands placed on individual stimuli. In Problem 1, Stimulus A was previously reinforced but now needed to be inhibited in Problem 3. Similarly, Stimulus C, which was not rewarded in Problem 2, became the correct choice in Problem 3. An elemental strategy would struggle to reconcile these conflicting contingencies, resulting in interference and degradation of earlier-learned associations.

In the second half of Stage 3 (50-100% of trials), this interference pattern remained. Performance on Problem 1 and Problem 2 was still not correlated (β = -0.598, *p* = 0.381), while performance on Problem 2 and Problem 3 was negatively correlated (β = -0.975, p = 0.004). No significant relationship was observed between Problem 1 and Problem 3 (β = 0.414, *p* = 0.616). The consistent interference between Problems 2 and 3 likely stemmed from the conflicting roles of Stimulus C. The absence of a clear relationship between Problems 1 and 3 may indicate that Stimulus A had become too unstable to support consistent performance due to its changing reward associations across contexts.

Taken together, these results suggest that marmosets relied on an elemental stimulus-response association strategy that is fundamentally unsuited to the relational demands of the TP task. As overlapping contingencies accumulated, interference increased, ultimately leading to widespread performance decline across all three problems.

#### Marmoset Transverse Patterning performance is unaffected by age

In humans, performance on the TP task declines with age, a deficit attributed to deterioration in hippocampal-dependent relational memory (Clark et al., 2017; Cohen et al., 1999; Ryan et al., 2016). In contrast, elemental learning, which relies on striatal systems, tends to remain stable across the lifespan (Bartus et al., 1979; Cox et al., 2008; Hill et al., 2023; Rapp, 1990). Given our findings that marmosets rely on an elemental strategy to approach the TP task, we asked whether aging would similarly affect TP performance in this species.

To address this question, we compared accuracy between young (< 8 years old) and aged (>= 8 years old) marmosets across each of the three Stages of the TP task. A two-way ANOVA revealed a significant main effect of Stage (Fig 2A, *F*(2, 66) = 7.08, *p* = 0.002), but no main effect of Age Group (*F*(1, 66) = 0.00, *p* = 0.99), and no Stage × Age Group interaction (*F*(2, 66) = 0.16, *p* = 0.86). Post hoc comparisons confirmed that young and aged marmosets performed similarly at every Stage (all *p* > 0.67). The main effect of Stage was driven by a significant decline of performance from Stage 2 to Stage 3 (*p* = 0.001), reflecting the group-level performance collapse already described, but this was not modulated by age. Treating age as a continuous variable yielded the same conclusion. There were no significant associations between age and accuracy in any stage (Fig 2B, Stage 1: β = 0.002, *p* = 0.933, Stage 2: β = 0.004, *p* = 0.513, Stage 3: β = -0.002, *p* = 0.545), further supporting the conclusion that transverse patterning performance in marmosets is not affected by age.

**Figure 2.**
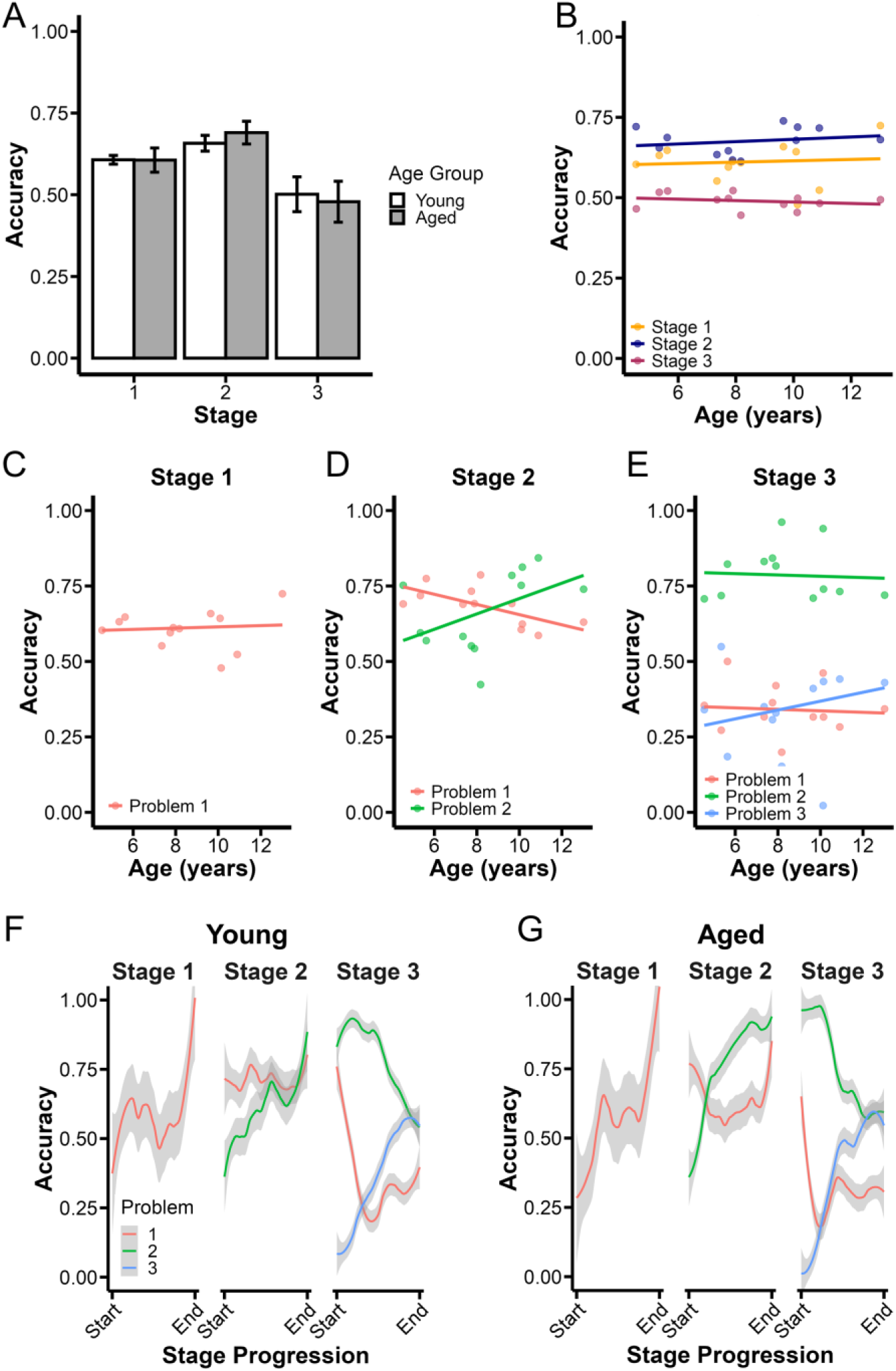
Transverse Patterning performance is unaffected by age in marmosets. (A) Mean accuracy ± SEM across stages of the TP task in Young (white bars) and Aged (gray bars) marmosets. There were no significant differences in performance between age groups on any stage. (B) Accuracy by age for all three stages. Each dot represents an individual marmoset. Regression lines show no consistent relationship between age and performance averaged across problems in any stage. (C-E) Stage-specific regression analyses showed no significant associations between age and performance in Stage 1 (C), Stage 2 (D), or Stage 3 (E). (F-G) LOESS-smoothed learning trajectories separated by age group. Young (F) and aged (G) marmosets show similar patterns of performance across all three stages, indicating that TP performance is not modulated by age.

To further investigate whether age influenced learning at a finer level, we conducted robust regression analyses predicting accuracy from age separately for each Problem within each Stage. In Stage 1, performance on Problem 1 was unrelated to age (Fig 2C, β = 0.002, *p* = 0.922), suggesting that initial acquisition was unaffected. In Stage 2, age was negatively associated with performance on Problem 1 (Fig 2D, β = -0.017, *p* = 0.003), but positively associated with Problem 2 (β = 0.025, *p* = 0.073). These opposite-sign effects were modest and difficult to interpret mechanistically. They may reflect variability during the transition to managing two problems rather than a meaningful age-related trend. In Stage 3, where all three overlapping problems were presented together, no significant age and performance associations were observed for any problem (Fig 2E, Problem 1: β = -0.002, *p* = 0.742; Problem 2: β = -0.002, *p* = 0.828; Problem 3: β = 0.015, *p* = 0.327).

Because age-related differences in strategy could theoretically drive subtle differences in performance, we also examined LOESS-smoothed learning trajectories separately for young (Fig 2F) and aged marmosets (Fig 2G). Both groups showed highly similar patterns: stable acquisition during Stages 1 and 2, followed by marked decline in Stage 3. This parallelism suggests that young and aged marmosets approached the TP task similarly, likely relying on a common elemental learning strategy that cannot support relational problem-solving.

Together, these findings suggest that, unlike in humans, aging does not impair TP performance in marmosets. This is consistent with the interpretation that marmosets rely on an elemental learning strategy, which remains stable across the lifespan and is insufficient to meet the relational demands of the TP task.

### Experiment 2

#### Marmosets successfully learn and retain multiple independent associations in a Concurrent Discrimination (CD) task

To determine whether marmosets’ failure on the TP task reflected a broader limitation in managing three problems simultaneously, we tested them on a Concurrent Discrimination (CD) task. The structure matched that of the TP task, three problems introduced across three stages, but unlike TP, the stimulus pairs in this task were non-overlapping, eliminating the need for relational processing.

Performance was analyzed using LOESS smoothing to visualize accuracy over time (Fig 3A). In Stage 1, all marmosets rapidly learned Problem 1, reaching high levels of accuracy. In Stage 2, they successfully learned Problem 2 while maintaining strong performance on Problem 1, indicating that learning a second problem did not interfere with retention of the first. In Stage 3, marmosets learned Problem 3 while continuing to perform well on Problems 1 and 2. These results demonstrate that marmosets can successfully learn and retain multiple stimulus-response associations concurrently, as long as those associations do not involve overlapping or conflicting structure and can therefore be learned independently without relational interference.

**Figure 3.**
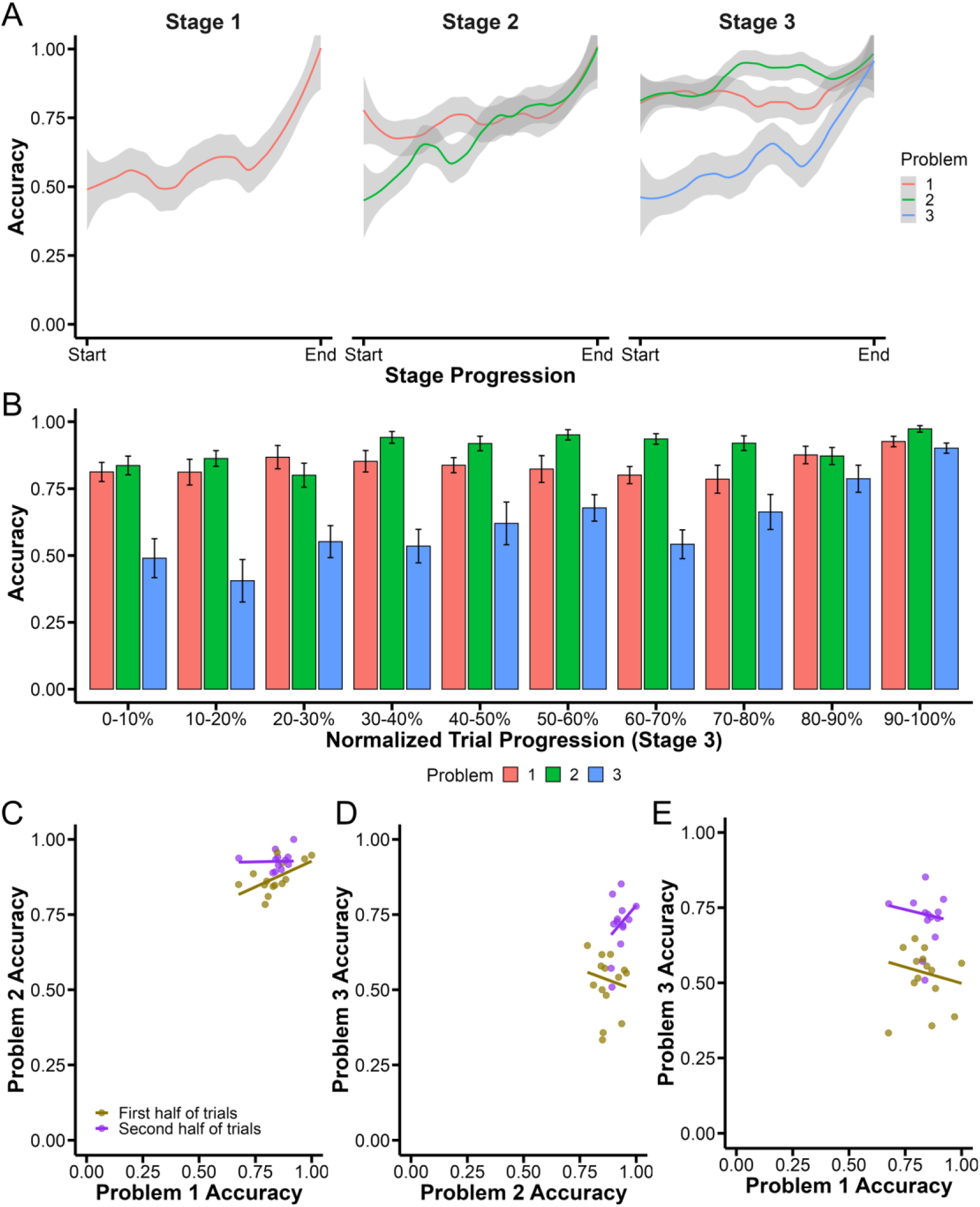
Marmosets successfully acquire multiple independent associations in the Concurrent Discrimination (CD) task. (A) LOESS-smoothed learning curves for of the three problems across each stage of the task. Accuracy steadily improves and criterion is achieved for all problems with no evidence of performance collapse in Stage 3. (B) Binned accuracy across Stage 3, shown in 10% increments of normalized trial progression. Marmosets maintained high accuracy on Problems 1 and 2 while gradually acquiring Problem 3, indicating successful learning of all three non-overlapping discriminations. (C-E) Accuracy correlations between problems in Stage 3. (C) Performance on Problems 1 and 2 was positively correlated during the first half of Stage 3 but not during the second half. (D, E) Problem 3 accuracy was not significantly associated with performance on either Problem 1 or 2, consistent with independent acquisition of non-overlapping associations and minimal interference.

To determine whether marmosets’ success on CD reflects an ability to manage multiple problems specifically when relational interference is absent, a key contrast with their failure on TP, we binned trials into 10% intervals and examined learning trajectories across problems (Fig 3B). Accuracy was analyzed using a two-way ANOVA with factors of Problem and Trial Bin. The analysis revealed a significant main effect of Problem (*F*(2, 390) = 161.19, *p* < 0.001), a significant main effect of Trial Bin (*F*(9, 390) = 10.39, *p* < 0.001), and a significant Problem x Trial Bin interaction (*F*(18, 390) = 4.32, *p* < 0.001), indicating that accuracy varied across problems and changed over time.

Post hoc comparisons revealed that in the early phase of Stage 3 (0-30% of trials), accuracy on Problem 3, introduced at that stage, was significantly lower than on Problems 1 and 2 (all *p*s < 0.001), consistent with its novelty. Performance on Problems 1 and 2, which were acquired in earlier stages, did not differ significantly across all bins (*p*s > 0.40), suggesting strong retention of previously learned associations. By the 40-70% bins, accuracy on Problem 3 had improved, but performance on this Problem remained lower than Problem 1 and 2 (*p*s < 0.05). Finally, by the 80–100% bins, accuracy across all three problems had fully converged, with no significant differences between them (all *p* > 0.20).

We next evaluated whether performance on each problem was correlated with each other during Stage 3. In the first half of Stage 3, performance on Problem 1 and Problem 2 was positively correlated, indicating that individuals who performed well on one problem also tended to perform well on the other (Fig 3C, *p* = 0.004). These associations were already learned during Stages 1 and 2, so this relationship likely reflects individual differences in overall learning ability, that is, animals who performed well on one early learned problem also tended to perform well on the other, not evidence of any meaningful integration across problems.

In contrast, performance on Problem 3, which was newly introduced in Stage 3, showed no significant correlation with performance on either Problem 1 or Problem 2 during either half of Stage 3 (Fig 3D-E, all *p*s > 0.15). This pattern suggests that marmosets acquired each problem independently, without linking new associations to prior ones. Such performance is consistent with an elemental learning strategy, in which each stimulus-response mapping is learned in isolation, rather than integrated through configural learning.

#### Aging does not affect learning of independent discriminations

We next asked whether aging affected marmosets’ ability to learn and retain multiple problems when those problems do not involve overlapping or conflicting elements. In humans, performance on tasks that rely on elemental learning remains stable across lifespan. If marmosets were using an elemental learning strategy to solve the CD task, we would expect performance to be age-invariant.

To test this, we conducted a two-way ANOVA with factors of Stage and Age Group (young vs aged). We found no main effect of Age Group (Fig 4A, *F*(1, 78) = 0.0397, *p* = 0.53) and no Stage x Age Group interaction (*F*(2, 78) = 0.39, *p* = 0.68), indicating that young and aged marmosets performed similarly across all stages. There was, however, a significant main effect of Stage (*F*(2, 78) = 13.57, *p* < 0.001), showing that performance improved as animals progressed through the task. Post hoc tests confirmed this improvement, with significant increases in accuracy from Stage 1 to Stage 2 (*p* = 0.008) and from Stage 1 to Stage 3 (*p* < 0.0001), and a trend toward continued gains from Stage 2 to Stage 3 (*p* = 0.0848). To complement this analysis, we also examined age as a continuous variable. Robust regression analyses showed no significant relationship between age and performance in any stage (Fig 4B, Stage 1: β = 0.008, *p* = 0.173; Stage 2: β = 0.004, *p* = 0.521; Stage 3: β = 0.002, *p* = 0.576).

**Figure 4.**
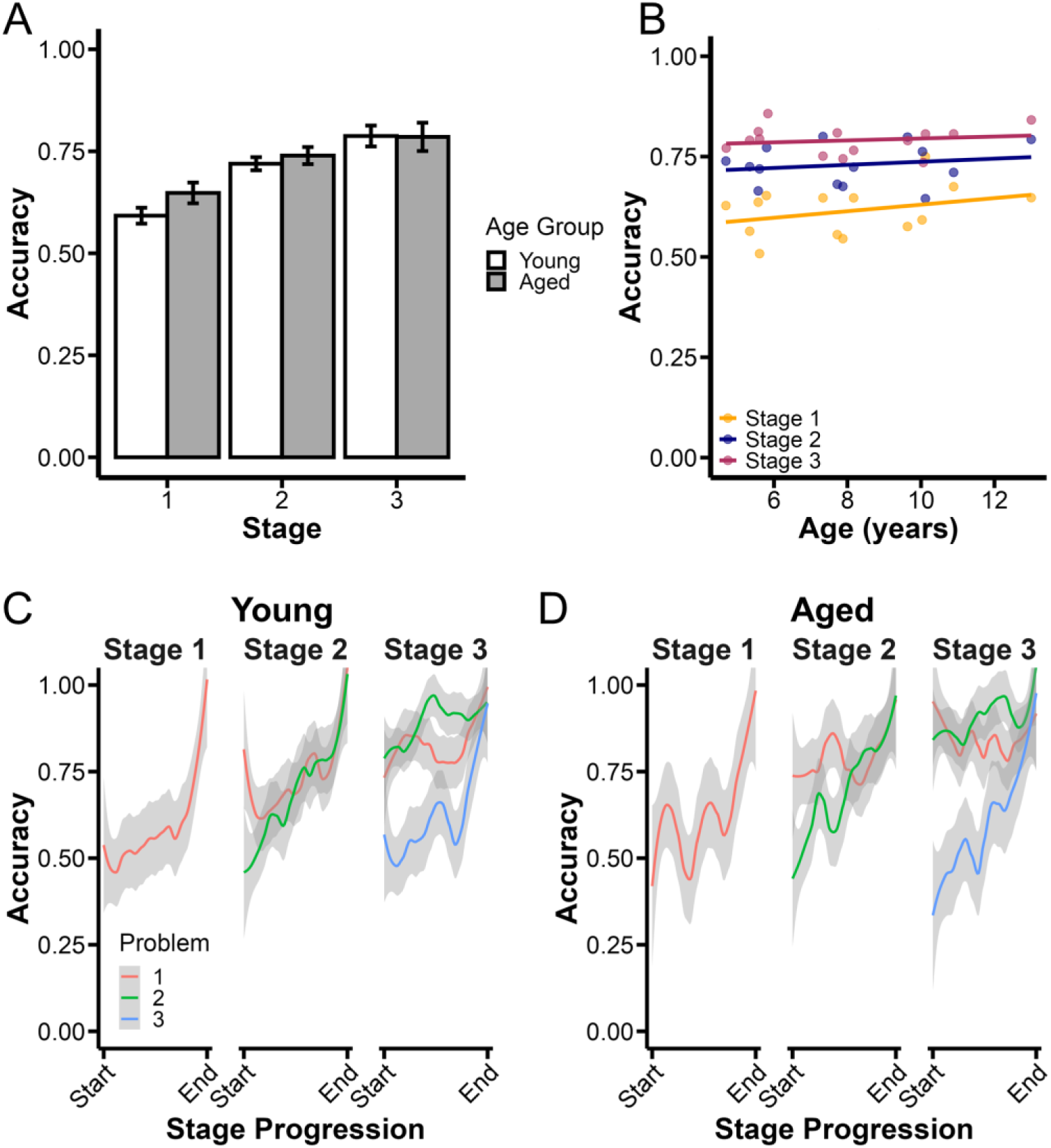
Age does not affect performance on the Concurrent Discrimination (CD) task. (A) Mean accuracy ± SEM for young and aged marmosets across each stage of the CD task. There were no differences between age groups. (B) Accuracy by age across all three stages. Each dot represents an individual marmoset. Regression lines indicate no significant association between age and performance averaged across problems in any stage. (C-D) LOESS-smoothed learning curves for each problem across stages, separated by age group. Young (C) and aged (D) marmosets showed similar acquisition patterns, demonstrating that aging does not impair the ability to learn or retain multiple independent discriminations.

Since age-related differences in strategy could potentially drive subtle differences in performance, we also examined LOESS-smoothed learning trajectories separately for young (Fig 4C) and aged marmosets (Fig 4D). Both groups showed highly similar patterns: stable acquisition during Stages 1, 2, and Stage 3 suggesting that young and aged marmosets approached the task similarly.

Taken together, these results show that aging did not impair the ability to learn or retain multiple discriminations when the problems could be solved independently. This is consistent with the idea that marmosets use an elemental learning strategy to solve this task, which remains stable across the adult lifespan.

### Experiment 3

#### Simultaneous introduction of all problems does not rescue transverse patterning performance

One possible explanation for marmosets’ failure on the TP task is that extended exposure to the first two problems (A+ B-and B+ C-) enabled animals to succeed using simple, stimulus-bound rules (e.g., “choose A if present, otherwise choose B”) rather than integrating the full relational structure. This may have prevented animals from integrating the third problem (C+ A-) when it was introduced in Stage 3. To test this hypothesis, we developed the Simultaneous Transverse Patterning (Sim-TP) task, in which all three problems were introduced simultaneously at the outset, eliminating the opportunity to build performance through isolated, non-relational rules.

Performance was analyzed using LOESS smoothing, which suggested that accuracy remained near chance levels across all three problems for the entire testing period (Fig 5A). To quantify this, we binned trials into 10% increments and conducted a two-way ANOVA with factors of Problem and Trial Bin. There were no significant main effects of Problem (Fig 5D, *F*(2, 330) = 0.015, *p* = 0.985), Trial Bin (*F*(9, 330) = 0.589, *p* = 0.806), or their interaction (*F*(18, 330) = 0.670, *p* = 0.840). These results indicate that marmosets failed to learn the relational structure of the TP task, even when all associations were introduced concurrently (Table 3).

**Figure 5.**
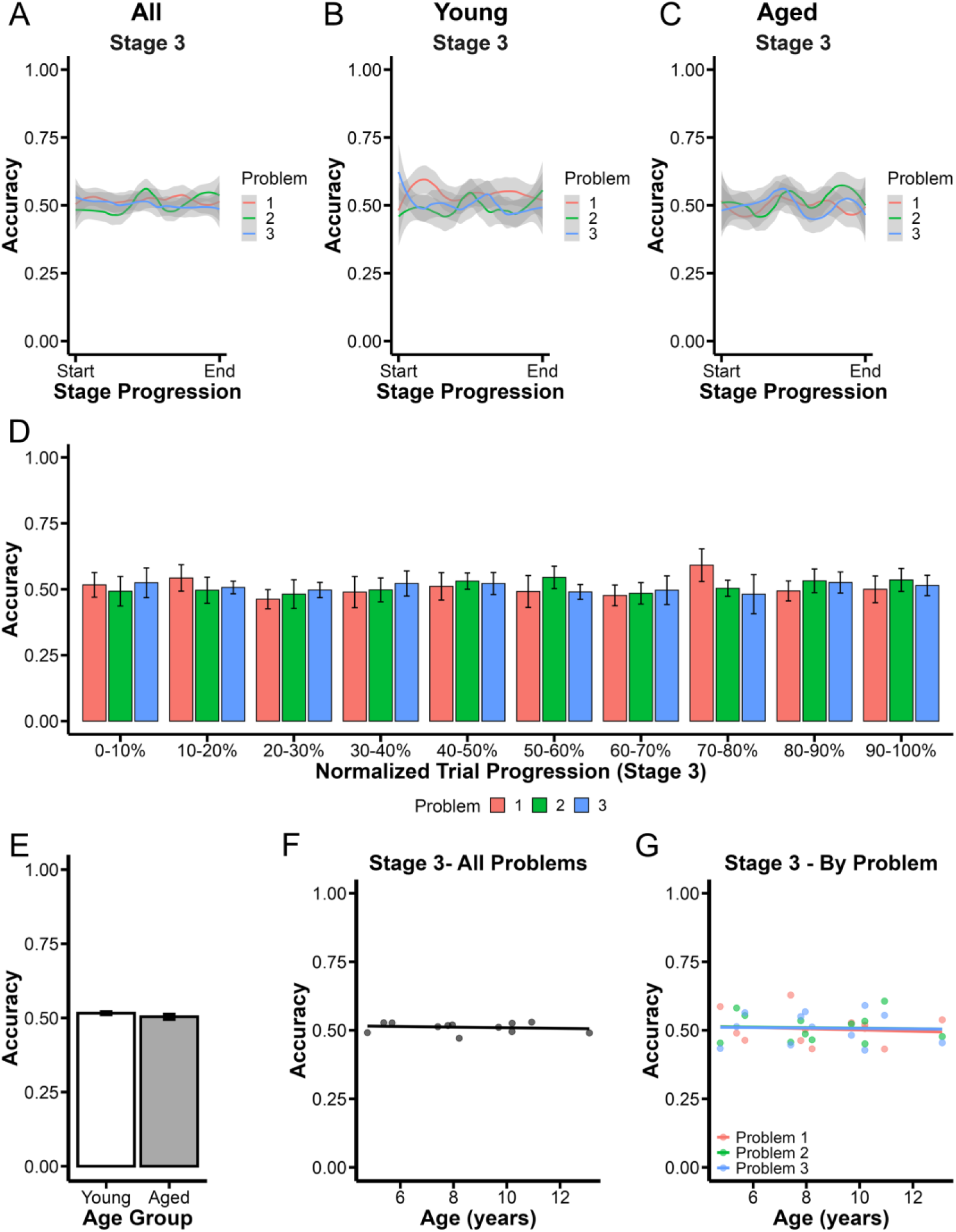
Marmosets fail to acquire the relational structure in the Simultaneous Transverse Patterning (Sim-TP) task, regardless of age. (A-C) LOESS-smoothed learning curves show accuracy across Stage 3 for all animals (A), young (B), and aged (C) marmosets. Accuracy remained near chance levels across all three problems indicating failure to acquire the overlapping associations. (D) Binned accuracy (10% increments) across Stage 3 confirmed no evidence of improvement over time on any problem. (E) Mean accuracy ± SEM across all three problems did not differ between young and aged marmosets. (F) Average accuracy on all problems across Stage 3 plotted by age showed no relationship between age and performance. (G) Accuracy for each problem in Stage 3 plotted by age confirmed that none showed age-related effects.

This finding does not indicate that extended exposure to earlier problems was the primary cause of failure on the TP task. Instead, it supports the conclusion that marmosets struggle with the TP task because of its relational demands and the interference that arises when elemental strategies are applied to overlapping contingencies.

#### Marmoset Transverse Patterning performance is unaffected by age

We next asked whether aging influenced performance on this Sim-TP version of the task. LOESS-smoothed trajectories showed that accuracy remained near chance for all three problems across the entire testing period for both young (Fig 5B) and aged (Fig 5C) marmosets. An independent samples *t*-test revealed no significant difference in accuracy between age groups (Fig 5E, *t*(8.1) = 1.146, *p* = 0.284). Treating age as a continuous variable also revealed no association between age and overall performance (Fig 5F, β = -0.001, *p* = 0.640), and no significant relationship between age and accuracy on any individual problem (Fig 5G, Problem 1: β = -0.002, *p* = 0.753; Problem 2: β = -0.001, *p* = 0.903; Problem 3: β = -0.001, *p* = 0.923). These results further support the idea that marmosets do not spontaneously engage relational memory systems, regardless of age or task structure, and instead rely on strategies that are ineffective for solving relational problems.

### Experiment 4

#### Marmosets can simultaneously learn multiple non-overlapping discriminations

To determine whether marmosets’ failure on the TP task reflects a general limitation in learning multiple problems simultaneously, or whether it’s driven specifically by interference from overlapping stimulus elements, we tested marmosets on a modified concurrent discrimination task. In this version, named Simultaneous Concurrent Discrimination (Sim-CD), all three problems were introduced simultaneously, but each used distinct stimuli, eliminating the need for relational memory.

LOESS-smoothed learning curves revealed that performance steadily improved on all three problems, with marmosets ultimately reaching criterion (Fig 6A; Table 3). To quantify this performance over time, we binned trials into 10% increments and ran a two-way ANOVA with factors of Problem and Trial Bin. The analysis revealed significant main effects of Time Bin (Fig 6D, *F*(9, 330) = 21.85, *p* < 0.001) and Problem (*F*(2, 330) = 4.39, *p* = 0.0131), as well as a Problem x Time Bin interaction (*F*(18, 330) = 1.78, *p* = 0.0265). Post hoc comparisons showed that during early training (0-30% of trials), performance was lower on Problem 3 compared to the others (*p*s < 0.05), likely reflecting its slower acquisition. However, these differences diminished over time, and by the second half of the task, there were no significant differences between problems, indicating that marmosets were able to learn and maintain all three discriminations in parallel when interference was removed.

**Figure 6.**
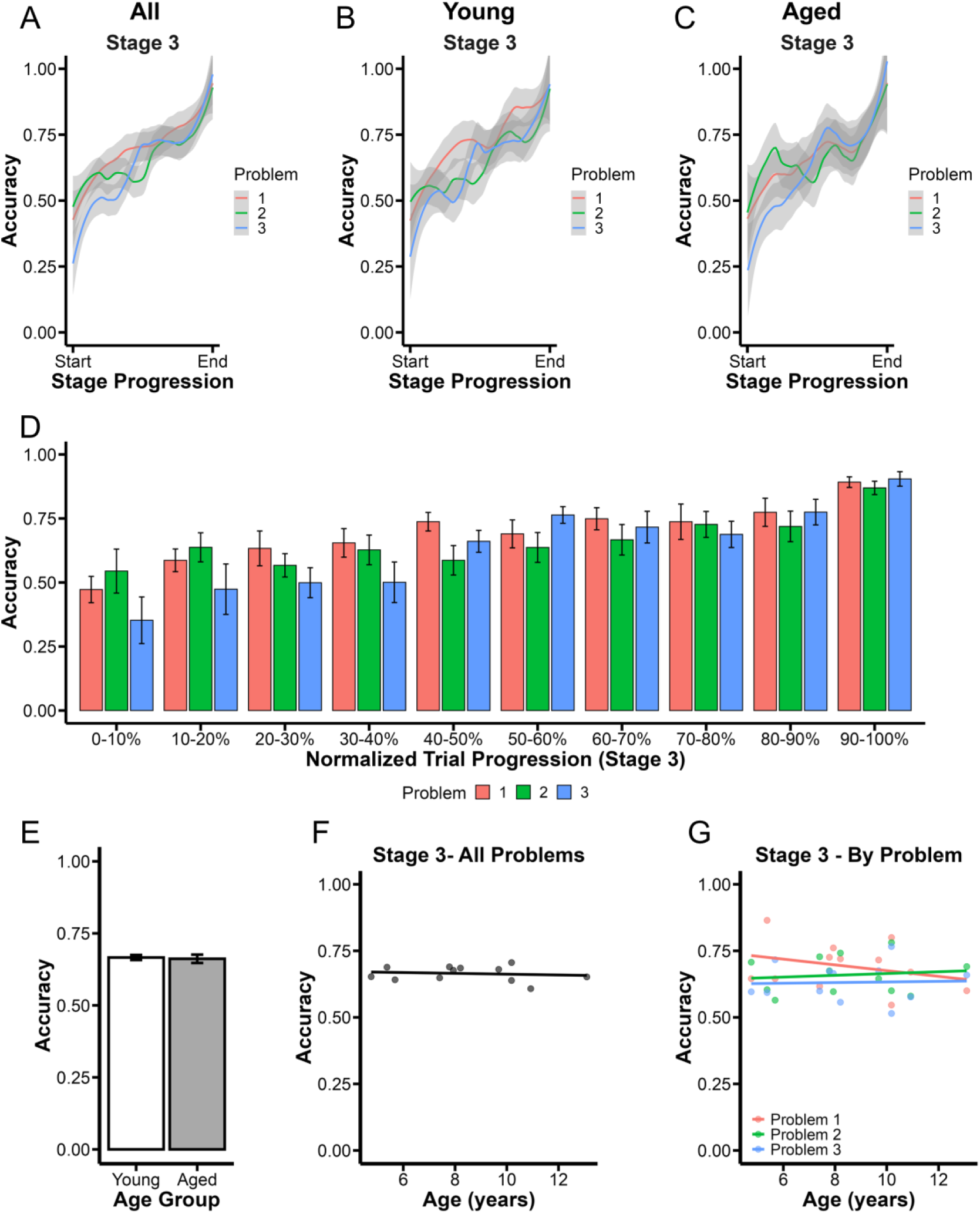
Marmosets successfully acquire multiple non-overlapping discriminations in the Simultaneous Concurrent Discrimination (Sim-CD) task, regardless of age. (A-C) LOESS-smoothed learning curves across Stage 3 show steady improvement across all three problems for all animals (A), young (B), and aged (C) marmosets, indicating robust acquisition of non-overlapping associations when all problems were introduced simultaneously. (D) Accuracy binned into 10% increments across Stage 3 reveals consistent improvement throughout training. (E) Mean accuracy ± SEM across all three problems did not differ between young and aged marmosets. (F) Average accuracy on all problems across Stage 3 plotted by age showed no relationship between age and performance. (G) Accuracy for each problem in Stage 3 plotted by age confirmed that none showed age-related effects.

#### Aging does not affect simultaneous learning of independent discriminations

We next asked whether age influenced performance on this Sim-CD task. We used LOESS smoothed learning curves to visualize learning curves in young (Fig 6B) and aged (Fig 6C) marmosets and observed qualitatively similar patterns. An unpaired *t*-test comparing overall accuracy between young and aged marmosets revealed no significant difference (Fig 6E, *t*(8.2) = 0.27, *p* = 0.794). Treating age as a continuous variable also showed no relationship between age and overall performance (Fig 6F, β = -0.002, *p* = 0.674). Lastly, to examine whether age might affect specific problems, we conducted separate regressions for each one. Again, no significant associations emerged (Fig 6G, Problem 1: β = -0.011, *p* = 0.352; Problem 2: β = 0.003, *p* = 0.679; Problem 3: β = 0.001, *p* = 0.887).

Together, these results demonstrate that marmosets are capable of simultaneously learning and maintaining multiple discriminations when the problems involve independent, non-overlapping stimuli. This contrasts sharply with their failure on the TP and Sim-TP tasks and strongly suggests that their difficulty with those tasks is a direct result of relational conflict between overlapping stimulus problems.

## DISCUSSION

Despite its widespread use as a benchmark for hippocampal function, the TP task produces strikingly inconsistent results across species. In humans, TP is highly sensitive to hippocampal damage and cognitive aging, reinforcing its status as a robust probe of relational memory (Astur & Sutherland, 1998; Rickard & Grafman, 1998). However, in non-human primates, outcomes have been mixed: some studies report impairments after hippocampal lesions, while others find no deficits or even improved performance (Alvarado & Bachevalier, 2005; Saksida et al., 2006). We addressed this question by testing common marmosets, a New World primate species increasingly used in cognitive neuroscience and aging research, on the TP task and three matched control tasks designed to dissociate elemental and configural learning strategies. Our results demonstrate that marmosets consistently fail TP when successful performance requires integration across overlapping stimulus relationships but readily succeed on structurally similar tasks that lack this relational demand. These findings suggest that, by default, marmosets approach TP using an elemental learning strategy based on simple stimulus-response associations, rather than flexibly engaging hippocampal-dependent configural learning. This insight helps resolve long-standing cross-species discrepancies and also underscores the importance of identifying the cognitive strategies animals use to solve a given task, particularly when interpreting performance in the context of aging or brain damage.

### Evidence for elemental learning in non-human primates

Marmosets failed to learn the TP task in both its standard form (Experiment 1; TP) and a version where all problems were introduced simultaneously to eliminate the possibility that extended training on early problems fostered reliance on simple, non-relational strategies (Experiment 3; Sim-TP). In both cases, performance deteriorated as interference from overlapping contingencies accumulated. In contrast, marmosets readily solved matched concurrent discrimination tasks in which stimulus overlap was removed, whether problems were introduced sequentially (Experiment 2; CD) or simultaneously (Experiment 4; Sim-CD). These findings rule out both a general inability to learn multiple problems and strategy bias introduced by task structure as explanations for TP failure. Instead, they implicate interference between overlapping associations as the key limiting factor.

The Complementary Learning Systems (CLS) model (McClelland et al., 1995; Norman, 2010) provides a framework for interpreting the dissociation between marmoset performance on TP and CD tasks. According to CLS, learning is supported by two parallel systems: the hippocampus, which rapidly forms pattern-separated representations to reduce interference, and cortical or striatal systems, which support slower, incremental learning of individual stimulus-response associations. The TP task, with its overlapping contingencies and hierarchical relationships (i.e., A > B > C > A), requires configural learning and is thus highly dependent on hippocampal processing. In contrast, the CD tasks, where each stimulus is uniquely paired and there is no overlap, can be solved through elemental learning strategies supported by striatal or cortical mechanisms.

The consistent failure of marmosets on TP, despite success on matched elemental tasks, suggests that they default to elemental learning, acquiring individual stimulus-response associations without integrating across problems. This strategy is effective in CD but breaks down in TP, where overlapping contingencies directly compete for associative strength. These findings align with prior work suggesting that non-human primates often rely on habit-based learning systems to solve discrimination tasks (Squire, 2004; Teng et al., 2000). Our results extend this literature by showing that marmosets persist with an elemental approach even when it is insufficient for task success, pointing to a strong bias toward elemental learning in marmosets.

While the evidence strongly favors an elemental learning approach, we considered the alternative possibility that marmosets attempted a configural strategy but failed to implement it successfully. To adjudicate between these interpretations, we examined behavior dynamically across trials and stages, focusing on how performance evolved over time rather than relying solely on static summary metrics. This trial-by-trial approach allowed us to capture subtle shifts in accuracy and interference across problems, providing a more nuanced view of the underlying learning strategies. We found that marmosets initially performed well on Problems 1 and 2, but performance collapsed following the introduction of Problem 3. Notably, this disruption was not uniform: performance on Problem 1 dropped below chance, whereas Problem 2 remained stable for a time before later declining. Such asymmetry is difficult to reconcile with a configural strategy, which should support stable performance once relational structure is encoded. Instead, the observed pattern reflects the kind of interference expected under an elemental framework, where overlapping contingencies directly compete for associative strength. The circular reinforcement structure of Stage 3 (A > B > C > A) poses a specific challenge to elemental learning, which lacks the flexibility to resolve mutually conflicting associations. Further supporting this interpretation, we observed negative correlations between problems during Stage 3, initially between Problem 3 and the others, and later between Problems 1 and 2, suggesting direct competition between stimulus–response mappings rather than integration into a unified relational structure. While we cannot definitively rule out failed configural learning as the TP failure mechanism, the overall behavioral dynamics strongly support reliance on elemental learning mechanisms that are ill-equipped to manage relational interference.

### Alignment with prior work in macaques

Our results also clarify longstanding questions in the macaque literature, highlighting how learning strategy and task design shape TP performance. In one study, macaques with hippocampal lesions outperformed intact controls on Stage 3 of TP, reaching high levels of accuracy while controls plateaued around 70% even after extensive training (Saksida et al., 2006). By contrast, humans reach criterion on Stage 3 rapidly, typically committing just a few errors. This apparent paradox, improved performance following hippocampal damage in macaques but impairment in humans, may reflect species differences in default learning strategies. Within the CLS framework, hippocampal and cortical systems operate in parallel but are tuned to different computational demands. If macaques primarily rely on striatocortical systems for habit-based learning, hippocampal removal may reduce interference from competing configural based representations, unmasking a more efficient elemental strategy. Humans, in contrast, approach TP with a hippocampal-dependent relational strategy, and thus are impaired when this system is compromised.

Interestingly, another study showed that hippocampal lesions impair TP performance in macaques, particularly under task conditions that minimize interference (Alvarado & Bachevalier, 2005). That study presented problems in blocks rather than interleaved them, allowing incremental acquisition of each problem in isolation and reducing the need to manage overlapping associations. Blocking may shift task demands to favor configural or mnemonic strategies, especially when remembering specific problem-reward contingencies is beneficial. Interleaving, by contrast, encourages generalization across problems and requires flexible updating, conditions that robustly engage the hippocampus. These procedural differences likely account for divergent lesion outcomes across studies and underscore the critical role of task structure in shaping the cognitive strategies and neural systems recruited across species.

### No age-related impairments on Transverse Patterning in marmosets

Aging is associated with well-documented structural and functional changes in the hippocampus, which are thought to underlie age-related deficits in memory (Radhakrishnan et al., 2022; Small et al., 2002; Stark & Stark, 2017; Yassa et al., 2011). In humans, aging impairs performance on the TP task, particularly in Stage 3, where all stimulus pairs must be integrated into a coherent relational structure (Driscoll et al., 2003; Gracian et al., 2016). This impairment is widely attributed to age-related hippocampal decline and supports the idea that TP performance in humans depends on hippocampal-dependent relational strategies (Driscoll et al., 2003).

In contrast, the influence of aging on TP performance in non-human primates has not been systematically explored. Here, we directly compared young and aged marmosets and found no significant age-related differences across any stage of the task. Both age groups followed similar learning trajectories consisting of initial improvement on Problems 1 and 2, followed by a decline in Stage 3, indicating that aging did not disrupt task acquisition or performance.

These findings are consistent with the interpretation that marmosets approach the TP task using habit-based strategies mediated by the striatum, rather than engaging hippocampal-dependent relational learning. Unlike the hippocampus, the striatum shows relatively preserved function with age, and prior studies have reported minimal age-related deficits in simple discrimination tasks that tap into striatal learning systems in non-human primates (Burke et al., 2010; Cox et al., 2008; Vanderlip et al., 2023). Our results extend this work by showing that even on a task classically used to probe hippocampal function in humans, marmosets show no age-related impairment, likely because they approach the task using neural systems that remain intact with age.

This species difference in age sensitivity further supports the idea that marmosets default to elemental learning strategies and highlights the importance of considering not just *what* task is being used in aging research, but *how* it is being solved. Strategy use plays a critical role in shaping task performance and its sensitivity to age, underscoring the importance of identifying how tasks are solved, not just whether they are, when evaluating their relevance for translational aging research.

### Ruling out alternative explanations for Transverse Patterning failure

While our earlier results strongly support the interpretation that marmosets approach the TP task using a habit-based strategy, we also systematically tested and ruled out several alternative explanations for their Stage 3 failure. One possibility is that marmosets are simply unable to form or maintain three distinct stimulus-response associations. To address this, we implemented a concurrent discrimination (CD) task with the same reward contingencies as TP but with non-overlapping stimuli. All marmosets successfully acquired the three problems, and in Stage 3, they maintained high accuracy on Problems 1 and 2 while improving on Problem 3, demonstrating the capacity to learn and retain multiple associations concurrently when interference is minimized.

Another possibility is that the standard TP task structure, in which Problems 1 and 2 are trained to criterion before Problem 3 is introduced, may have encouraged the use of simple, non-relational strategies. With extensive exposure to the first two problems (A+ B- and B+ C-), animals may have developed simple non-relational rules (e.g., “choose A if present, otherwise choose B”) that allowed for successful performance without engaging hippocampal-dependent relational processing. These rules could support strong performance on the early problems where there is a linear hierarchical relationship among stimuli, but would break down once overlapping contingencies accumulate with the introduction of Problem 3. To evaluate this, we tested a Sim-TP condition in which all three problems were introduced simultaneously, eliminating the opportunity to build simple, stimulus-bound strategies based on extended exposure to a subset of problems. If inflexibility were the limiting factor, performance should have improved. Instead, marmosets performed at chance across all three problems for the duration of training, suggesting that equal exposure does not mitigate the interference.

To further rule out general difficulty with simultaneous learning, we repeated the simultaneous presentation format using non-overlapping problems, as in the CD task. Under these conditions, marmosets once again learned all three associations without difficulty. Finally, performance on the control tasks did not differ between young and aged marmosets, suggesting that aging did not alter the learning strategy used. This further supports the idea that marmosets consistently relied on the same habit-based approach across all conditions.

Together, these control experiments provide strong converging evidence that the marmosets’ failure in Stage 3 of the TP task is not due to an inability to learn multiple associations, nor to the sequential introduction of problems or simultaneous problem presentation. Rather, the critical limiting factor is interference between overlapping associations, a challenge that habit-based systems are ill-equipped to resolve due to the absence of pattern separation mechanisms. This pattern of results aligns with a learning system dominated by elemental encoding and highlights the need for hippocampal engagement to successfully disambiguate relational structure.

### Implications

This study offers critical insights into how cognitive tasks are solved across species, with direct implications for translational neuroscience. Our results show that marmosets fail the Transverse Patterning task not because of cognitive decline or limited memory capacity, but because they approach the task using a fundamentally different learning strategy. Whereas humans rely on hippocampal-dependent relational processing to integrate overlapping information, marmosets appear to default to an elemental, habit-based strategy supported by corticostriatal systems. This strategy allows them to acquire individual stimulus-response associations but leaves them vulnerable to interference when relational structure is required.

These findings challenge the assumption that differences in task performance reflect deficits, rather than species-specific differences in cognitive strategies and neural systems engaged during task execution. Importantly, they highlight the need to validate that a task engages the intended neural system before using it to assess function, particularly in translational models of aging or neurodegenerative disease. A task that reliably probes hippocampal integrity in humans may not do so in marmosets, depending on how the task is approached cognitively. Without this validation step, researchers risk misinterpreting behavioral impairments as evidence of neural dysfunction when the task may not actually engage the targeted brain system.

More broadly, this work underscores the importance of aligning not just task structure, but also cognitive demands and strategy use across species. Behavioral homology requires more than surface-level similarity; it requires that tasks tap into equivalent computations and neural mechanisms. By identifying the learning strategies marmosets use in a task traditionally used to probe hippocampal function, this study provides a critical foundation for refining cognitive assays in non-human primates. Doing so will improve the interpretability and translational relevance of animal models in research on aging, neurodegeneration, and brain injury.

## Conclusion

Together, our results demonstrate that marmosets approach the Transverse Patterning task using an elemental, habit-based learning strategy rather than the relational, hippocampal-dependent system engaged by humans. This strategy enables the acquisition of simple stimulus-response associations but fails when contingencies overlap and require flexible integration. Through a series of carefully designed control experiments, we ruled out alternative explanations including limited memory capacity, the effects of sequential problem introduction, and difficulty learning multiple problems simultaneously, demonstrating that the core limitation stems from interference between overlapping associations, a challenge that habit-based systems are ill-equipped to resolve.

These findings underscore the importance of identifying not just whether a task is learned, but how it is learned, as cognitive strategies and neural systems they engage can differ dramatically across species, even when the task structure is held constant. Without careful validation of the processes underlying task performance, researchers risk drawing incorrect conclusions about cognitive capacity or neural decline. By clarifying the learning mechanisms marmosets use in a classic hippocampal task, this study provides essential groundwork for developing species-appropriate cognitive assays and for improving the interpretability of behavioral outcomes in studies of aging, memory, and neurodegeneration.

## Acknowledgments

This research was supported by an AHA-Allen Initiative in Brain Health and Cognitive Impairment award made jointly through the American Heart Association and The Paul G. Allen Frontiers Group: 19PABH134610000AHA, National Institutes of Health grant 1R21AG068967-01, grants from the Larry L. Hillblom Foundation and the Don and Lorraine Freeberg Foundation, and the Fiona and Sanjay Jha Chair in Neuroscience. We thank Katie Williams for assistance in the care of the marmosets and technical support.

## Notes

### Competing Interest Statement

The authors have declared no competing interest.

